# Characterizing chemotherapy-induced neutropenia and monocytopenia through mathematical modelling

**DOI:** 10.1101/2020.04.02.022046

**Authors:** Tyler Cassidy, Antony R. Humphries, Morgan Craig, Michael C. Mackey

## Abstract

In spite of the recent focus on the development of novel targeted drugs to treat cancer, cytotoxic chemotherapy remains the standard treatment for the vast majority of patients. Unfortunately, chemotherapy is associated with high hematopoietic toxicity that may limit its efficacy. We have previously established potential strategies to mitigate chemotherapy-induced neutropenia (a lack of circulating neutrophils) using a mechanistic model of granulopoiesis to predict the interactions defining the neutrophil response to chemotherapy and to define optimal strategies for concurrent chemotherapy/prophylactic granulocyte colony-stimulating factor (G-CSF). Here, we extend our analyses to include monocyte production by constructing and parameterizing a model of monocytopoiesis. Using data for neutrophil and monocyte concentrations during chemotherapy in a large cohort of childhood acute lymphoblastic leukemia patients, we leveraged our model to determine the relationship between the monocyte and neutrophil nadirs during cyclic chemotherapy. We show that monocytopenia precedes neutropenia by 3 days, and rationalize the use of G-CSF during chemotherapy by establishing that the onset of monocytopenia can be used as a clinical marker for G-CSF dosing post-chemotherapy. This work therefore has important clinical applications as a comprehensive approach to understanding the relationship between monocyte and neutrophils after cyclic chemotherapy with or without G-CSF support.

## 1 Introduction

In humans, approximately 100,000 hematopoietic stem cells (HSCs) produce 100 billion blood cells a day (Lee-Six et al., 2018). This astonishing hyper-productivity accounts for the hematopoietic system being one of the most intensively studied and best understood stem cell systems (Mackey, 2001). All terminal blood cells originate from HSCs that differentiate, proliferate, and mature along many lineages before reaching the circulation to perform their multitude of functions, including oxygenating the body (red blood cells), clotting and wound repair (platelets), and providing immunity from intruders (white blood cells). Neutrophils are important immune regulators and key components of the innate immune response produced by HSCs differentiating into the myeloid lineage, and subsequently undergoing a period of exponential expansion and maturation before transiting into the circulation following sequestration in the bone marrow reservoir (Craig et al., 2016). Monocytes are also myeloid cells that mature within the bone marrow from monoblasts before exiting to the circulation as monocytes, where they remain for several days before marginating in the tissues (Swirski et al., 2014). Tissue-resident monocytes differentiate into macrophages and dendritic cells as a crucial component of an effective immune response (Pittet et al., 2014).

All blood cell production and function is orchestrated through the interaction of a class of small proteins with receptors on the surfaces of HSCs, progenitors, and differentiated cells. These proteins, known as cytokines and chemokines, are produced either by the blood cells themselves or by organs such as the kidneys, liver, and spleen (Roberts, 2005; Qian et al., 1998; Athanassakis and Iconomidou, 1995). Cytokines act to regulate blood cell production, whereas chemokines direct blood cells through chemotaxis. Though cytokines have overlapping functions to ensure the robustness of the hematopoietic system, many are primary modulators of a specific lineage.

Granulocyte-colony stimulating factor (G-CSF) is the principal cytokine responsible for the regulation of neutrophil production and function (Craig et al., 2015). Its main roles are to modulate the egress of cells from the bone marrow reservoir into the circulation, and control the speed of maturation, proliferation, and differentiation of HSCs into the neutrophil lineage (Craig, 2017). G-CSF and neutrophils are inversely related to each other, i.e. when the number of neutrophils in the circulation decreases, G-CSF concentrations rise to stimulate their release and production and vice versa.

This cytokine paradigm (that concentrations of circulating blood cells are reciprocally related to their main cytokine regulator, i.e. a negative feedback) is echoed throughout the hematopoietic system. In particular, monocyte production and function are principally controlled by granulocyte-macrophage colony-stimulating factor (GM-CSF) and macrophage stimulating factor (M-CSF), which act similarly to how G-CSF acts on the neutrophils (Rapoport et al., 1992) to produce monocytes from the HSC, regulate their differentiation into macrophages, and mediate their role in the immune response.

Disrupted cytokine networks are involved in a host of pathologies, including rare oscillatory hematological diseases like cyclic neutropenia (Dale and Hammond, 1988; Dale and Mackey, 2015; El Ouriaghli et al., 2003) and cyclic thrombocytopenia (Krieger et al., 2018; Langlois et al., 2017), atherosclerosis (Nahrendorf et al., 2010), and anemia (Mackey, 1978). Neutropenia and monocytopenia are characterized by a lack of neutrophils or monocytes, respectively, in circulation and have important consequences for the host’s ability to mount an effective immune response. Some cytopenias are inherited and result from mutations to cytokine receptors that inhibit the proper association of cytokines to receptors. Others may be acquired through transitory events including the action of drugs, including cytotoxic agents.

Cytotoxic chemotherapy is designed to disrupt cellular division to control cancer growth. Unfortunately, this cytotoxicity can have significant deleterious effects on other dividing cells in the body, including the rapidly-renewing terminally-differentiated neutrophils (Dale and Mackey, 2015). Chemotherapy-induced neutropenia and ensuing infections are a common side effect of cytotoxic chemotherapy (Brooks et al., 2012; Craig et al., 2015; Friberg et al., 2002; Glisovic et al., 2018), necessitating dose size reductions or complete therapy cessation. These therapy modifications leave the patient particularly vulnerable to infection, a major cause of treatment-related mortality even in malignancies with high survival rates (Gatineau-Sailliant et al., 2019; Glisovic et al., 2018). Thus, there is intense interest in dampening the adverse hematological events associated with cytotoxic chemotherapy, through the design of less toxic chemotherapy combinations or the introduction of prophylactic agents. To that end, G-CSF is frequently administered as a rescue drug (prophylactically or as an adjuvant) during cytotoxic chemotherapy (Brooks et al., 2012; Craig et al., 2015). The optimal scheduling of exogenous G-CSF during chemotherapy is an active area of research, and recent mathematical modelling efforts suggest that delaying G-CSF after chemotherapy reduces the incidence and severity of chemotherapy-induced neutropenia (Craig et al., 2015; Krinner et al., 2013; Vainas et al., 2012).

Mathematical modelling has long played a role in understanding healthy and pathological hematopoietic dynamics. Early models aimed to delineate HSC and hematopoietic dynamics (Glass and Mackey, 1979; Loeffler and Wichmann, 1980; Mackey, 1978, 1979; Wichmann and Loeffler, 1985; Wichmann et al., 1988) and understand the so-called ‘dynamic diseases’ (Mackey, 2020-submitted) like cyclic neutropenia (Haurie et al., 1999; von Schulthess et al., 1983) and thrombocytopenia (von Schulthess and Gessner, 1986). Since the discovery of cytokines and their control of hematopoiesis, there has been more recent attention to the role cytokines play for endogenous maintenance of basal cell concentrations (Craig et al., 2016) and on the use of exogenous administration of cytokine mimetics for treatment (Brooks et al., 2012; Craig et al., 2015; Foley and Mackey, 2009; Quartino et al., 2012, 2014; Schmitz et al., 1996; Câmara De Souza et al., 2018). Extensive reviews of mathematical models of hematopoiesis are available in Pujo-Menjouet (2016) and Craig (2017).

Despite the role of monocyte derived macrophages (MDMs) in the resolution of infection, monocyte production has not been extensively studied using mathematical models. To our knowledge, most mathematical modelling efforts that include MDMs have revolved around the innate immune response to infection (Álvarez et al., 2017; Day et al., 2009; Eftimie et al., 2016; Marino et al., 2015; Smith, 2011; Smith et al., 2013). As a particular example, Smith et al. (Smith, 2011) developed a mathematical model for the resolution of pneumonia infection in mice that included three distinct phases of the immune response to pneumonia, including the immune response from tissue resident macrophages and cytokine driven recruitment of circulating neutrophils, and the role of MDMs in infection clearance. There, recruitment of MDMs was driven by the neutrophil concentration in the infected area. Their model shows good agreement with murine data, and has been used to model pneumonia co-infection with influenza and the action of antibiotics (Smith et al., 2013; Schirm et al., 2016). Other formalisms, including hybrid ODE and agent based models, have also been used to study the role of MDMs in infection. For example, Marino et al. (Marino et al., 2015) studied the role of macrophage polarization in *M.tuberculosis* infection. While these models underscore the importance of MDMs in the resolution of infection, their focus was primarily on the site of infection and therefore did not describe the regulation of the production of neutrophils and monocytes in the bone marrow, assuming cytokine- or neutrophil-driven recruitment instead.

Conversely, the role of MDMs in the immune response to malignant tumours has been extensively studied (Boemo and Byrne, 2019; Eftimie and Eftimie, 2018; Mahlbacher et al., 2018; Owen et al., 2011). Modelling work has focused on macrophage interactions in hypoxic tumours, with recent work also studying the effects of macrophage polarization in tumour progression (Eftimie and Eftimie, 2018). Similar to the infection modelling discussed earlier, these models do not study the production of monocytes in the bone marrow, since they are principally focused on understanding tumour-immune dynamics in the tumour microenvironment and not descriptions of systemic monocytopoiesis.

Chemotherapy affects circulating monocyte concentrations and may influence tumour-macrophage interactions. Accordingly, understanding the dynamics of monocyte production during chemotherapy is crucial. Though chemotherapy-induced monocytopenia has previously been linked to an increased risk of developing neutropenia (Kondo et al., 1999), and GM-CSF can be incorporated as a rescue drug during cytotoxic chemotherapy (Tafuto et al., 1995), little attention has been paid in the mathematical literature to understanding monocytopenia during chemotherapy.

Here we address the issue of monocyte production during chemotherapy by developing a novel model of monocytopoiesis based on our previous work modelling granulopoiesis (Craig et al., 2016). This model is derived directly from the mechanisms of monocyte production and parameterized from homeostatic relationships and a limited amount of data fitting. Leveraging data from a cohort of childhood acute lymphoblastic leukemia patients, we then applied our combined model to understand monocyte and neutrophil dynamics during cyclic chemotherapy. We found that there is an important relationship between the observation of monocytopenia and ensuing neutrophil nadir during cytotoxic chemotherapy. Our results have potentially significant clinical implications as they suggest that the arrival of the monocyte nadir can be used as an indicator to mitigate neutropenia through exogenous G-CSF administration.

## 2 Neutrophil and monocyte modelling

### 2.1 Model of granulopoiesis

We have previously described a mathematical model for the physiology of neutrophil production that explicitly accounts for free and bound G-CSF concentrations (*G*_1_(*t*) and *G*_2_(*t*), respectively) and their pharmacodynamic effects (Craig et al., 2016). Briefly, the model accounts for the hematopoietic stem cell population *Q*(*t*), neutrophils sequestered in the bone marrow reservoir after undergoing exponential proliferation and a period of maturation (denoted by *N*_*R*_(*t*)), and neutrophils in circulation, *N*(*t*). The full set of model equations is given by

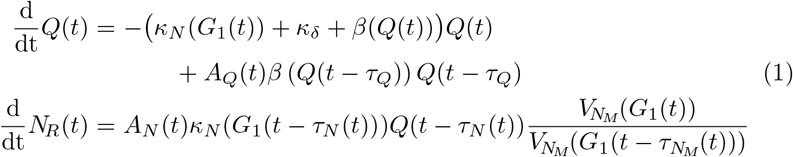

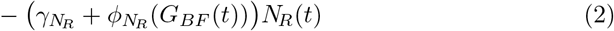

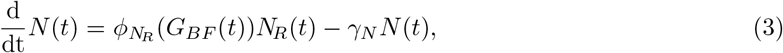

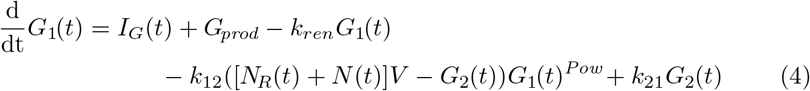

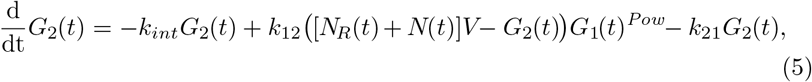

with parameter definitions and estimations as in Craig et al. (2016). Throughout, *κ*_*i*_, *A*_*i*_, and *γ*_*i*_ denote differentiation, amplification due to proliferation, and death rates, respectively, in the *i*-th lineage. Variations on this model have previously been applied to the study of cyclic neutropenia (Colijn and Mackey, 2005) and to optimize G-CSF regimens during chemotherapy (Brooks et al., 2012; Craig et al., 2015; Foley et al., 2006).

### 2.2 Pharmacokinetic and pharmacodynamic models

Cytotoxic chemotherapy agents interrupt cell division (through disruptions to microtubule assembly or DNA synthesis) to kill malignant cells. Contrary to modern targeted therapies, this broad cytotoxicity also affects rapidly-dividing hematopoietic progenitor cells in the bone marrow. We model circulating drug concentrations as impacting on granulopoiesis and monocytopoiesis (described below) by accounting for the anti-cancer agent’s pharmacokinetics (PK). These PKs then determine the chemotherapeutic drug’s pharmacodynamics (PD).

The PKs of chemotherapy are described by a four-compartment model (central *C*_*p*_, fast-exchange *C*_*f*_, and two slow-exchange compartments 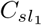 and 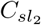) by

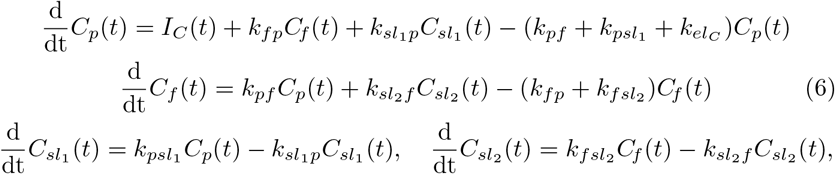

where *k*_*i*_ indicate rate parameters with values as in Craig et al. (2016).

Since neutrophils no longer divide after exponential expansion, we consider only the HSCs and progenitor cells to be affected by plasmatic chemotherapy concentrations. Monocytes can differentiate from classic (CD14+CD16−) to non-classic (CD16+) types in the blood (described below), but we discount the effects of chemotherapy on this conversion, owing to the short half-life of the drug in the circulation and the relatively low evolution rate from classic to non-classic type. Accordingly, the PD effects of chemotherapy are modelled as

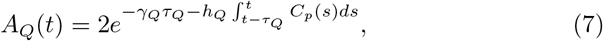

i.e. a decrease in the effective amplification (difference between proliferation and death over the time for HSC self-renewal) and by

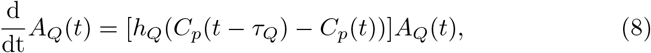

i.e. a decrease in the proliferation rate for neutrophil and monocyte progenitor cells. In both Eqs. (7) and (8), *τ*_*Q*_ represents the time for HSC self-renewal and *h*_*Q*_ denotes the effect of chemotherapy on HSC amplification *A*_*Q*_.

G-CSF is an endogenous cytokine that is administered as a drug during chemotherapy. Since the action of G-CSF is to bind to the surface of target cells, we model the pharmacokinetics of G-CSF using a two-compartment model accounting for free and bound concentrations (Eqs.(4) and (5), respectively) where *I*_*G*_(*t*) accounts for exogenous administration. As described in Craig et al. (2016), increasing G-CSF concentrations induce egress from the neutrophil bone marrow into circulation at rate 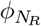, and act upstream to restock the mature reservoir by increasing the speed of maturation 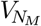, the rate of progenitor proliferation 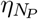, and the differentiation rate *κ*(*G*_1_) from the HSCs into the granulocyte lineage. These actions are modelled (from HSCs to circulation) as

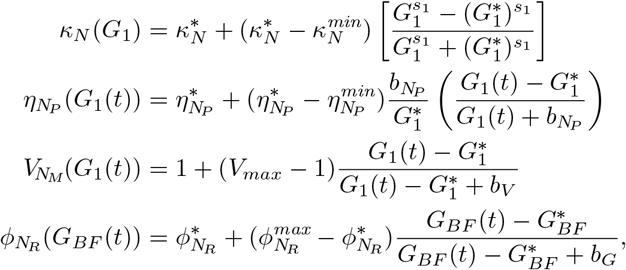

where *G*_*BF*_ denotes the fraction of G-CSF bound to neutrophil receptors, and all parameters have been estimated in Craig et al. (2016). Similar to the HSCs, we included the effect of cytotoxic chemotherapy on the proliferation of neutrophils by

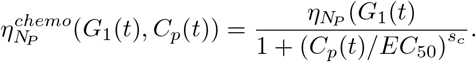

### 2.3 Mathematical model of monocyte production

We model monocytopoiesis similarly to our granulopoiesis model. A schematic of both granulopoiesis and monocytopoiesis is given in Fig. 1.

**Fig. 1.**
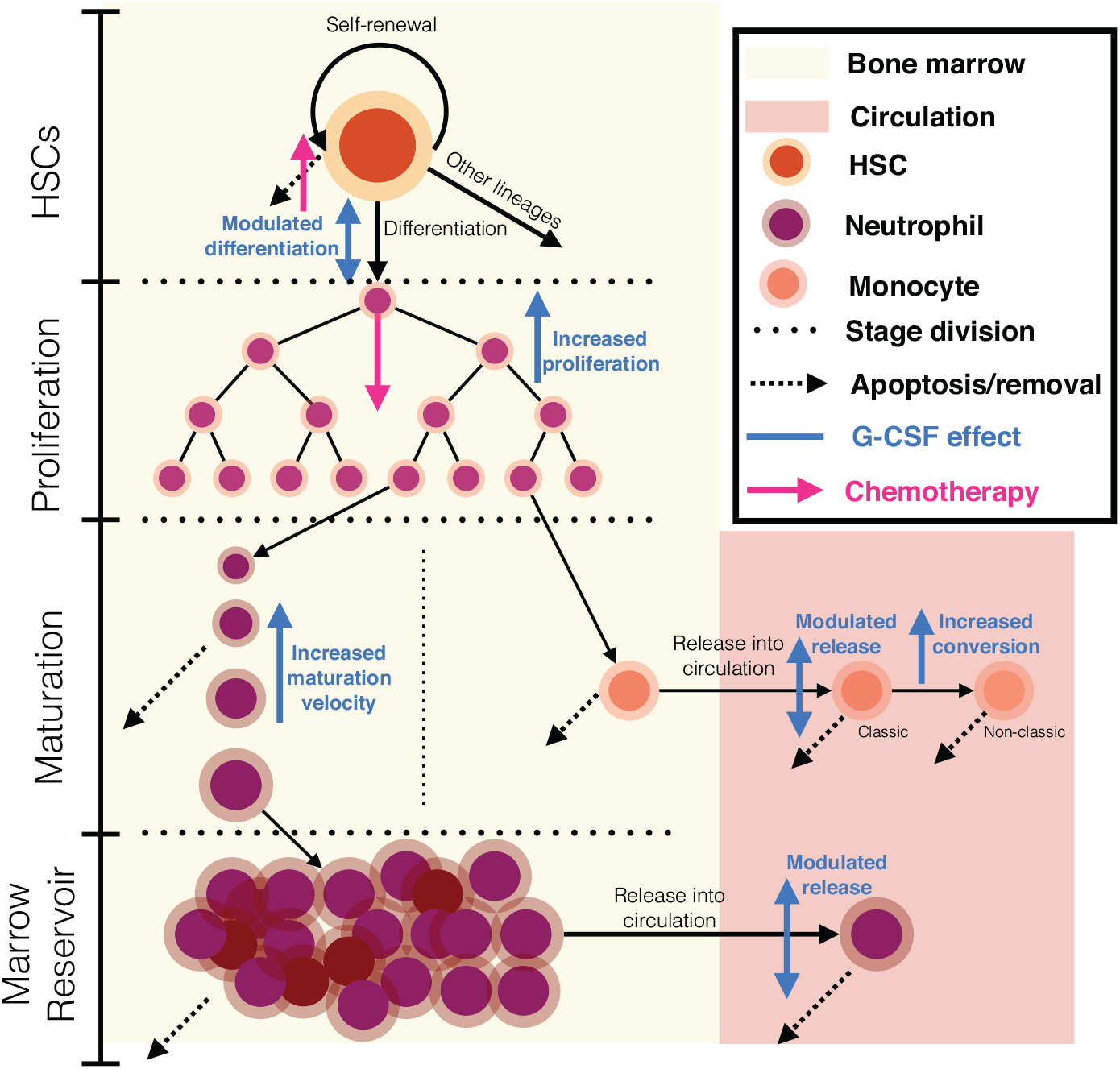
Production of neutrophils and monocytes. Hematopoietic stem cells self-renew to maintain their population, die, or differentiate into the myeloid (or other) lineages. Progenitors then undergo a period of exponential expansion before committing to becoming neutrophils or monocytes. Circulating neutrophils leave the bone marrow after a period of maturation and sequestration in the mature marrow reservoir. Monocytes are released into the circulation, where they may convert from classic to non-class subtypes. Both circulating neutrophils and monocytes are removed from the circulation through apoptosis.

#### 2.3.1 From stem cell to monocyte

As a simplifying modelling assumption, circulating G-CSF concentrations are used as a cipher for GM-CSF’s control of production and proliferation of monocyte precursors by all cytokines. The HSC differentiation rate into the monocyte lineage is given by *κ*_*M*_(*G*_1_(*t*)), implying that the flux of cells entering the monocyte lineage at a given time is *κ*_*M*_(*G*_1_(*t*))*Q*(*t*). The G-CSF dependent differentiation rate is

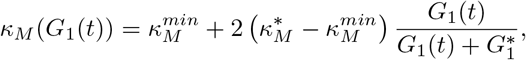

where the homeostatic concentration of unbound G-CSF is 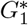. Equation (1) models differentiation from HSCs into the monocyte, platelet and erythrocyte lineages via *κ*_*δ*_. Therefore, we augment the differential equation for the HSCs to explicitly include differentiation into the monocyte lineage by

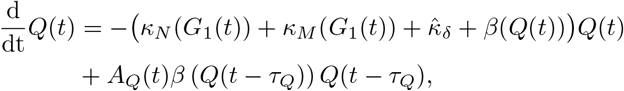

where 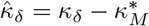.

After differentiation, monocyte precursors undergo both proliferation and death, with G-CSF acting to decrease the death rate of precursors. Rather than modelling these as separate processes, we combine them into an *effective* proliferation rate given by 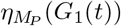, similar to the granulopoiesis model (Craig et al., 2016). Monocyte proliferation has been shown to persist even in the absence of G-CSF and other stimulating factors (Boettcher and Manz, 2017). Consequently, we model 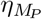 as a saturable and monotonically increasing function of circulating G-CSF with 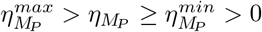, where

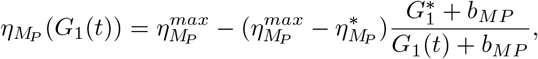

with 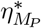 denoting the effective proliferation rate at homeostasis. This definition requires a constraint on *b*_*MP*_ to ensure that 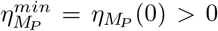. We model the effect of chemotherapy on proliferating monocytes similarly to the neutrophils via

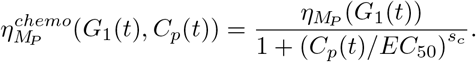

To model the proliferative process, we use an age structured partial differential equation (Craig et al., 2016; Cassidy et al., 2019; Metz and Diekmann, 1986) where the age variable models progression from HSC to mature monocyte. The density of monocyte precursors with age *a* at time *t* is denoted by *M*_*P*_(*t*, *a*). Then, as in the granulopoiesis model, the density of monocyte precursors satisfies the age structured partial differential equation (PDE)

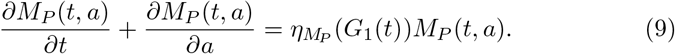

We assume that monocyte precursors leave the proliferative process in a deterministic process upon reaching the threshold age *a* = *a*_*M*_. To define an initial value problem for the age structured PDE (9), appropriate initial and boundary conditions must be provided. As mentioned, HSC-derived monocyte precursors enter directly into proliferation at age *a* = 0, which gives the boundary condition

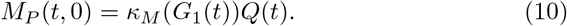

Analogously, the initial condition of (9) defines the initial density of monocyte precursors throughout the ageing process, and is given by

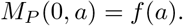

In general, the function *f*(*a*) is assumed to be integrable. However, as we will show, the biological interpretation of *M*_*P*_(*t*, *a*) provides a natural choice of initial density *f*(*a*), while for general *f* ∈ *L*_1_(0, ∞), the corresponding initial value problem admits a solution *M*_*P*_(*t*, *a*) (Perthame, 2007).

The density of monocyte precursors along the characteristic lines of (9) is

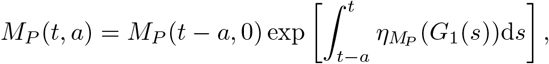

where

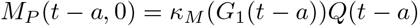

represents the flux of HSC into the monocyte lineage at time *t* − *a*, and the exponential term models population expansion due to proliferation.

Together, these relationships define the natural form for the initial condition *f*(*a*). The density of monocyte precursors at time *t* = 0 given by *f*(*a*) signifies that HSCs differentiated into the monocyte lineage some time in the past. Specifically, cells with age *a* at time 0 enter the monocyte lineage at time *t* = −*a*, when the flux of HSCs into the monocyte lineage is *κ*_*M*_(*G*_1_(−*a*))*Q*(−*a*). These monocyte precursors proliferate from their entrance into the lineage until time *t* = 0. Thus, the initial density *f*(*a*) is given by

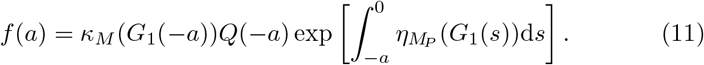

Similar to the neutrophil lineage, we model the time delay between differentiation into the granulocyte-monocyte lineage and the production of a mature monocyte by 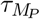. We assume for simplicity that the proliferative stage takes the same time for monocytes as neutrophils, so 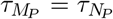. Moreover, we assume that monocyte precursors mature into classic monocytes after reaching the threshold age *a* = *a*_*m*_ over 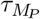 time units. Accordingly, the flux of mature classic monocytes entering the bone marrow is

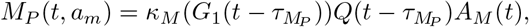

where

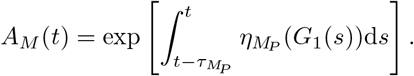

These nascent classic monocytes reside in the bone marrow before entering circulation (Nguyen et al., 2013; Mitchell et al., 2014; Van Furth et al., 1973; Mandl et al., 2014). As for the neutrophil reservoir in Craig et al. (2016), we model the bone marrow concentration of classic monocytes as

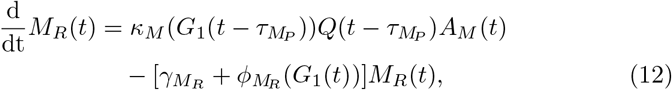

where 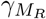 is the death rate of bone marrow monocytes and 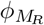 denotes the rate of release of classic monocytes into circulation.

Experimental data has shown an increase in circulating monocytes in response to G-CSF treatment (Hartung et al., 1995; Molineux et al., 1990), which we model by a G-CSF-dependent 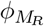. However, during viral infections, circulating G-CSF levels increase without a noticeable increase in monocytes (Pauksen et al., 1994). Thus, slightly increased G-CSF concentrations do not lead to monocyte release from the bone marrow, while administration of large amounts of G-CSF does induce depletion of the bone marrow monocyte concentration and a corresponding increase in circulating monocyte concentrations. To account for this threshold G-CSF concentration, we introduce 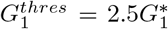 to denote the minimal G-CSF concentration inducing increased monocyte recruitment, with 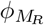 given by

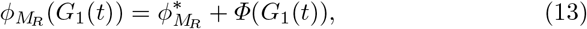

where

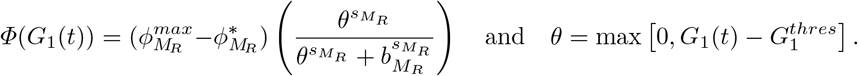

Here, 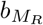 represents the G-CSF concentration for half maximal release of monocytes from the bone marrow. We assume that, if the G-CSF concentration is large enough to trigger increased monocyte egress from the bone marrow, then this release will be near maximal, rather than gradual, and thus set 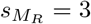.

#### 2.3.2 Monocyte dynamics in circulation

Egress of classic monocytes from the bone marrow into circulation is modulated by CCR2 (Patel et al., 2017; Shi and Pamer, 2011). Circulating classic monocytes are then either cleared from the circulation at a constant rate 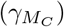 or evolve into non-classic monocytes (Patel et al., 2017). Classic and non-classic monocytes are distinguished by their expression of CD14 and CD16 surface markers. While circulating monocytes express CD16 in a continuous spectrum (Shantsil et al., 2011), we separate the monocyte population into classic (low CD16) and non-classic (high CD16) compartments.

Increases to the population of CD16+ monocytes have been observed during inflammatory conditions (West et al., 2012; Patel et al., 2017; Strauss-Ayali et al., 2007; Wong et al., 2011). We therefore model the evolution of classic monocytes into non-classic monocytes at the G-CSF-dependent rate *ν*(*G*_1_(*t*)) (Ingersoll et al., 2011; Kratofil et al., 2017; Wong et al., 2012), and assume that the evolution from classic to non-classic monocyte takes place at a saturable rate given by

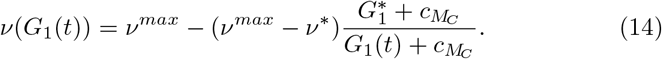

The circulating classic monocyte population dynamics are then governed modelled by

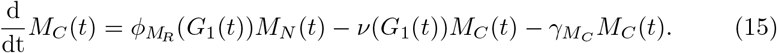

Since we assume that non-classic monocytes are only produced via evolution from the classic compartment and are cleared at a constant rate 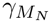, the dynamics of the non-classic monocytes are given by

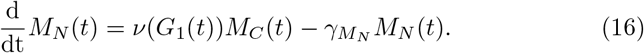

Taken together, the complete model of monocyte production is

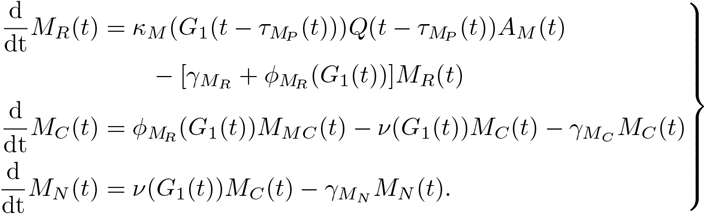

When coupled with Eqs. (1)-(5), the combined granulocytopoiesis-monocytopoiesis model is a system of state dependent discrete delay differential equations. To define the corresponding initial value problem, we must prescribe initial data defined over the delay interval. While we use homeostatic initial conditions throughout, this initial data typically is a continuous function defined over [*t*_0_ − *τ*, *t*_0_], where 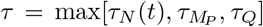 is the maximal delay. Here, *τ*_*N*_(*t*) is defined as in Craig et al. (2016) as the solution of the integral condition

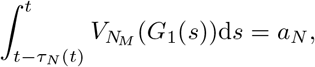

where 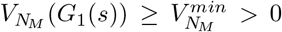, so 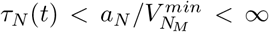 and *τ* is well-defined and finite. Accordingly, the appropriate phase space for the mathematical model is the infinite dimensional space *C*_0_([−*τ*, 0], ℝ^8^). Further, the results from Câmara De Souza et al. (2018) and Cassidy et al. (2019) imply that solutions of the mathematical model evolving from non-negative initial data remain non-negative.

## 3 Results

### 3.1 Monocyte parameter estimation at homeostasis

Monocytes and neutrophils share a significant portion of their early developmental pathway, and the point of divergence between the granulocyte and macrophage lineage is not well characterized. Accordingly, we assume that differentiation into the monocyte-macrophage lineage satisfies 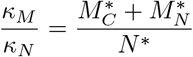, which gives 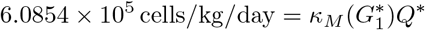.

The homeostatic circulating concentration of monocytes is *M*\* = *M*_*C*_ + *M*_*N*_ = (0.034 − 0.062) × 10^9^ cells/kg (Lichtman, 2016; Meuret et al., 1974; Meuret and Hoffmann, 1973; Whitelaw, 1972). In the calculations that follow, we assume that *M*\* = 0.060 × 10^9^ cells/kg and a 90% : 10% ratio of classic to non-classic monocytes (Patel et al., 2017; Strauss-Ayali et al., 2007; Wong et al., 2012, 2011; Ziegler-Heitbrock, 2014; Zimmermann et al., 2010). Thus, 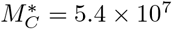 cells/kg and 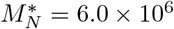 cells/kg.

Patel et al. (2017) measured the appearance of deuterium label in circulating monocytes following a 3-hr pulse labelling and concluded that monocytes spend between 1.5 and 1.7 days in the bone marrow following proliferation before entering circulation. This yields the average residence time in the bone marrow and the homeostatic transit rate from the bone marrow into circulation by setting

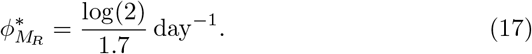

To calculate the concentration of mature monocytes, we assume that influx and efflux of classic monocytes are balanced at homeostasis, i.e.

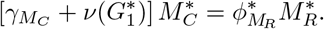

Classic monocytes have a circulating half-life of roughly *t*_1/2_ = 1 day (Ginhoux and Jung, 2014; Patel et al., 2017), which accounts for both clearance of classic monocytes from circulation and evolution into non-classic monocytes. Therefore,

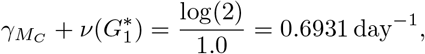

The bone marrow concentration of monocytes at homeostasis is therefore

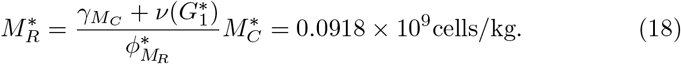

The total bone marrow monocyte pool (TBMMP) consists of monocyte precursors and mature monocytes. The exponential nature during proliferation provides a natural upper bound on the number of monocyte lineage cells in the bone marrow. Since DNA synthesis in eukaryotic cells takes approximately 8 hours, to compute this upper bound, we allow 1.5 hours for the remainder of the cell cycle and assume that cells cannot complete more that 2.5 divisions. This corresponds to a division time of 9.6 hours. Then, assuming that there is no death of proliferating cells at homeostasis, there are at most

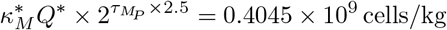

proliferating monocyte precursors. In our model, the number of monocyte precursors in the bone marrow at homeostasis is given by

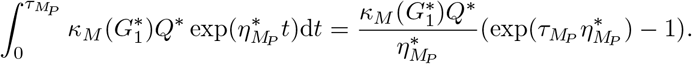

Moreover, there are 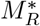 mature monocytes in the bone marrow at homeostasis. Therefore, the TBMMP is

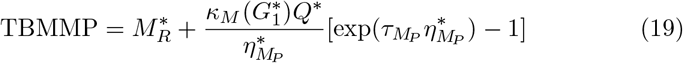

Re-arranging equation (19) gives

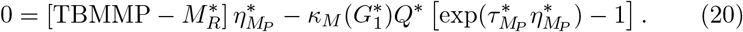

Then, for a given TBMMP, the only unknown in (20) is the homeostatic proliferation rate 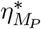 that must satisfy 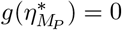, where

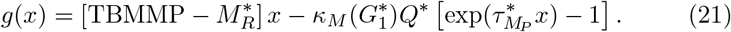

Since

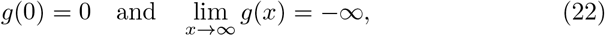

and *g*′′(*x*) < 0, there is a unique strictly positive solution 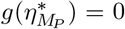 if *f*(*x*) is strictly increasing at 0. Further, *g*(*x*) is increasing at *x* = 0 if

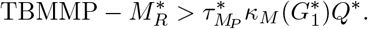

Accordingly, there can only be effective proliferation if the monocyte precursor pool is strictly larger than the accumulation of monocyte precursors solely due to input from the HSCs. With 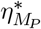 from Eq. (20), the death rate of bone marrow monocytes, 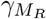, can then be calculated from

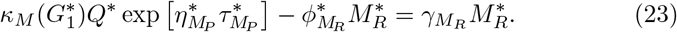

Non-classic monocytes spend roughly 5 − 7 days in circulation (Ginhoux and Jung, 2014; Patel et al., 2017), so

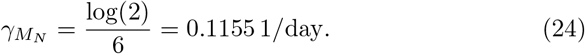

Therefore, at homeostasis,

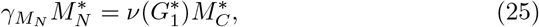

and

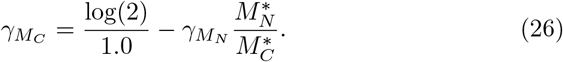

It has been observed during inflammatory conditions in trauma patients that non-classic monocytes comprise roughly 15% of the monocyte population on average (West et al., 2012). Thus, if 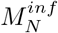 and 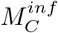 represent the inflammatory concentration of non-classic and classic monocytes, respectively,

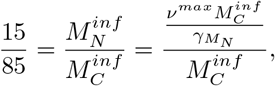

which gives a first approximation for 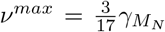. In the absence of any knowledge of the precise mechanism underlying cytokine driven evolution of classic to non-classic monocytes, we set 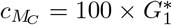.

Finally, TBMMP, 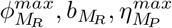 and 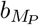 remain to be estimated to fully parameterize the monocytopoiesis model. With the exception of TBMMP, these parameters represent the response of cells in the monocyte lineage to increased G-CSF concentrations. Accordingly, we integrated data for circulating monocyte concentration following the administration of exogenous G-CSF in healthy humans to estimate these parameters by minimizing the *L*_2_ distance between our model’s prediction and a spline interpolating the average data reported in Hartung et al. (1995) (Fig. 2). To validate our estimates, we compared model predictions to *a new* data set of two sequential administrations of G-CSF (Hartung et al., 1995), and found that our model was able to accurately capture the monocyte dynamics in the second G-CSF administration cycle (Fig. 2).

**Fig. 2.**
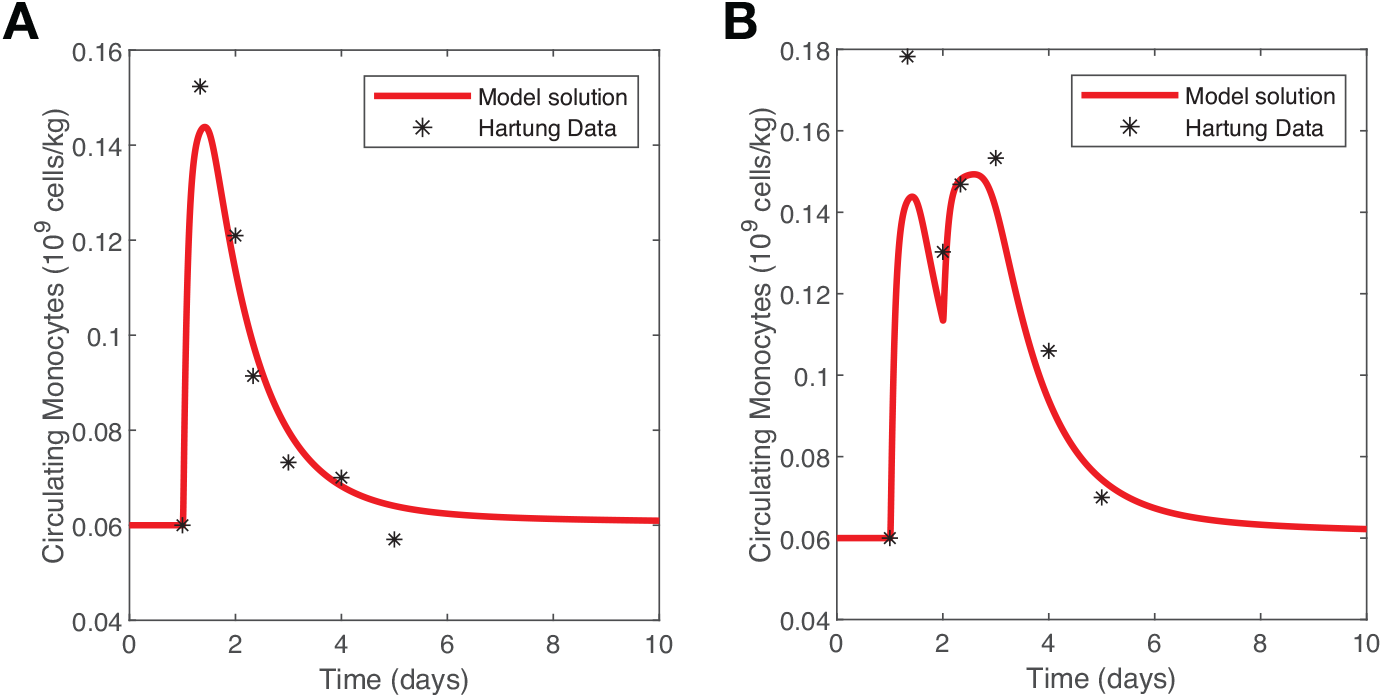
Parameter fitting results. A) Monocyte model fitting to the Hartung data for administration of a single dose of 480 000 *μ*g/kg G-CSF to healthy human volunteers. B) Monocyte prediction to Hartung data from administration of two doses of G-CSF to healthy volunteers.

Values for all parameters are listed in Table 1, and are comparable to previous measurements and estimates. For example, Van Furth et al. (1980) estimated that there are 0.1043 × 10^9^ cells/kg mature monocytes in the bone marrow, while 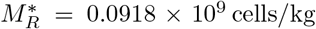. Whitelaw (1972) predicted an egress rate of 1.34 × 10^7^ cells/kg/day, while we calculate 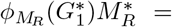 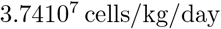. Our prediction that the evolution of classic into non-classic monocytes represents 1.8% of classic monocyte clearance is also close to the transition percentage of 1.4% calculated by Patel et al. (2017).

**Table 1.** Summary of parameter values as calculated in the text.

| Parameter | Value | Biological Interpretation | Source |
| --- | --- | --- | --- |
| TBMMP | $0.1619 \times 10^9$ cells/kg | Total bone marrow monocyte | Fit |
| $M_R^*$ | $0.0918 \times 10^9$ cells/kg | Bone marrow monocyte concentration | (18) |
| $\gamma_{MR}$ | 0.7654 /day | Bone marrow monocyte clearance rate | (23) |
| $\eta_{MP}^*$ | 1.5518 /day | Proliferation Rate | (20) |
| $\eta_{MP}^{max}$ | 1.5620 /day | Proliferation Rate | Fit |
| $b_{MP}$ | 1.1383 ng/mL | Proliferation half effect of G-CSF | Fit |
| $b_{MR}$ | 3.9810 ng/mL | Proliferation half effect of G-CSF | Fit |
| $\phi_{MR}(G_1^*)$ | 0.4077 /day | Release rate into circulation | (17) |
| $\gamma_{MC}$ | 0.6803 /day | Classic monocyte clearance rate | (26) |
| $\nu(G_1^*)$ | 0.0128 /day | Classic to non-classic evolution rate | (14) |
| $\gamma_{MN}$ | 0.1155 /day | Non-classic monocyte clearance rate | (24) |

### 3.2 Parameter sensitivity analysis

To assess the dependency of model predictions to parameter values, we performed a local sensitivity analysis of the monocytopoiesis model (see Craig et al. (2016) for discussion of parameter values in the neutrophil model). As in Cassidy and Craig (2019); Crivelli et al. (2012), we quantify the sensitivity of the model output to changes of ±10% in each parameter value. We further evaluate the time between the monocyte and neutrophil nadirs and the duration of monocytopenia as they are clinically-relevant. The results of this sensitivity analysis show that model predictions are robust to parameter variations (Fig. 3).

**Fig. 3.**
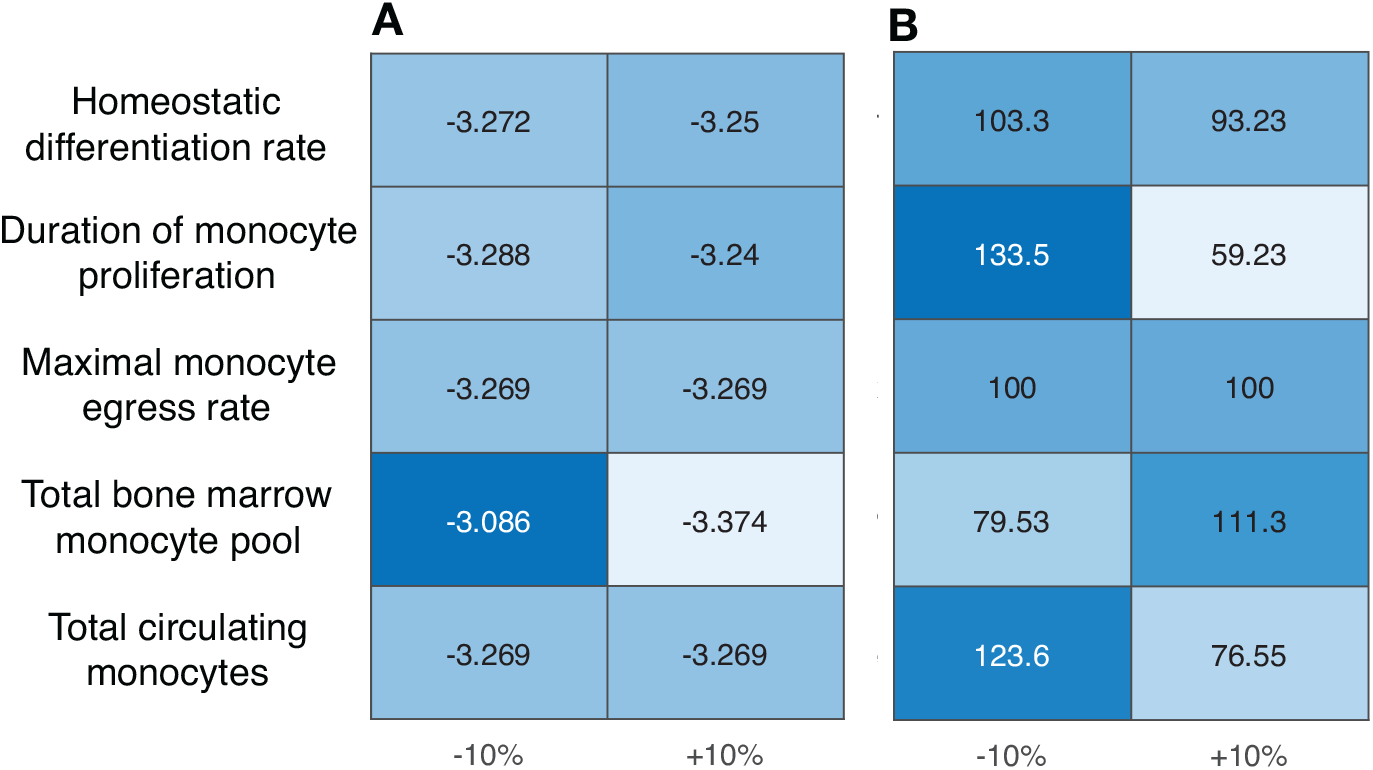
Results of local parameter sensitivity analysis. A) Time delay between the nadir of monocyte and neutrophil counts. The negative values indicates that the monocyte nadir occurs before the neutrophil nadir. B) Parameter sensitivity in duration of monocy-topenia when compared to the duration of monocytopenia with fitted parameters. In both cases, parameters were individually varied by 10% from their fitted values.

### 3.3 Monocytopenia precedes neutropenia

Acute lymphoblastic leukemia (ALL) is characterized by an overabundance of immature lymphocytes and is the most common childhood malignancy. Treatment differs defined based on a clinical risk-assessment, but childhood ALL is generally treated by chemotherapy administered in multiple phases depending on the protocol. Despite a high rate of recovery (five-year survival of 94% for children aged 0-14 in Canada (Canadian Cancer Statistics Advisory Committee, 2019)), pediatric patients frequently suffer from neutropenia and monocytopenia during treatment.

Kondo et al. (1999) found that monocyte counts can be predictive for ensuing grade 3 or 4 neutropenia during 3- or 4-week cyclic chemotherapy, which could be explained by discordant production times between monocytes and neutrophils. To assess the impact of cyclic chemotherapy on monocytopoiesis and granulopoiesis, we analyzed neutrophil and monocyte counts from 286 pediatric ALL patients treated with the Dana-Farber Cancer Institute (DFCI) ALL Consortium protocols DFCI 87-01, 91-01, 95-01, or 00-0120-22 at the Sainte-Justine University Health Center (Montreal, Canada). Previous genomic analysis of this cohort has revealed loci (DARC, GSDM, and CXCL2) predictive of neutropenic complications (Gatineau-Sailliant et al., 2019; Glisovic et al., 2018), as measured by the absolute phagocyte count or APC. We have also identified resonance (chemotherapy induced oscillations) in the neutrophil counts of 26% of these patients during their treatment (Mackey et al., 2020-submitted). Here, monocyte and neutrophil numbers were collected at 3-week intervals after induction, prior to the beginning of the next chemotherapy cycle.

We begin by performing classical statistical analyses (Pearson coefficient analysis) on the childhood ALL cohort to investigate possible mechanisms between monocyte counts and ensuing neutropenia during cyclic chemotherapy. Across all patients, we find a very low correlation (average Pearson correlation coefficient *r* = 0.27) between monocyte and neutrophil numbers (Fig. 4A). However, a positive correlative relationship between monocytes and neutrophils is detectable in a smaller subgroup (moderate positive Pearson correlation of *r* = 0.53; Fig. 4B).

**Fig. 4.**
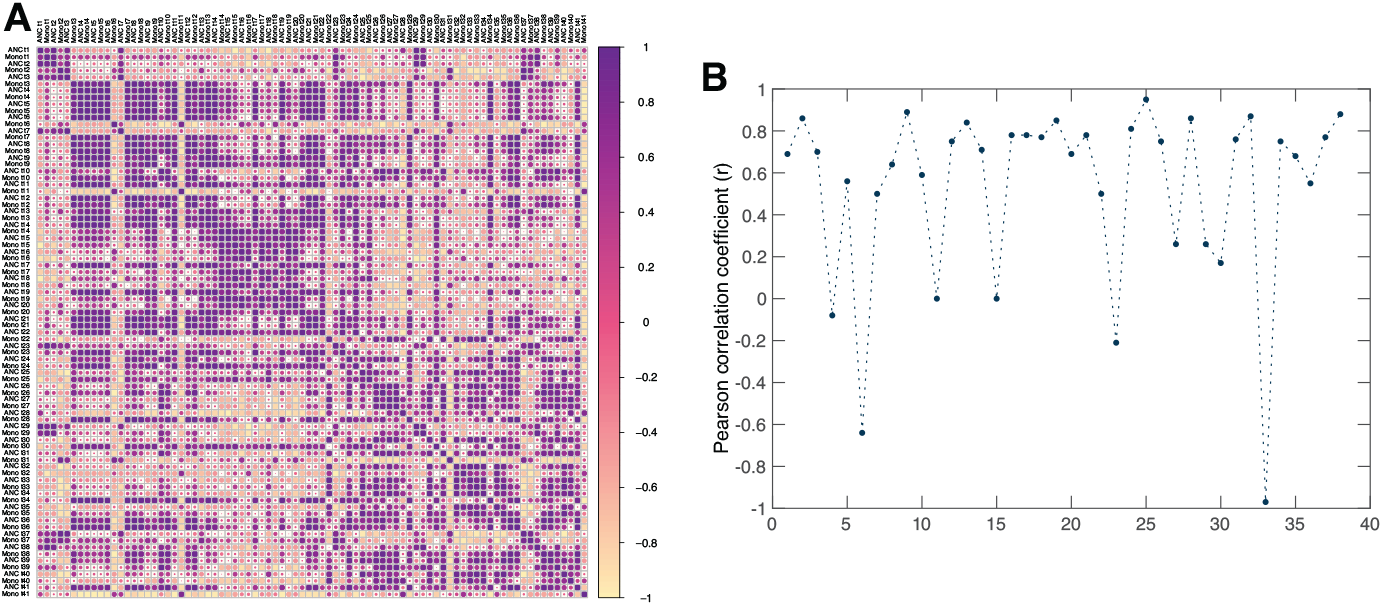
Correlations between monocyte and neutrophil counts in the childhood ALL cohort. A) Correlations between monocyte (Mono) and neutrophil (ANC) counts at each sampling point (*t*) in the full 286 patient cohort. Neutrophil and monocyte sampling points are indicated on horizontal and vertical axes. Deep purple circles represent perfect positive correlation i.e. that monocyte and neutrophil counts at sampling point *ti* and *tj* are positively predictive for one another. Deep yellow indicates a strongly negative correlation. As indicated by the lack of clustering, no discernable correlative relationship was found within the complete cohort. B) Restriction of A to comparison of monocytes to neutrophils at the same time point. At each sampling point, the Pearson correlation coefficient between monocytes and neutrophils is plotted. Though some points demonstrate a negative correlation, an average moderate positive correlation (r=0.53) was found, indicating that monocyte and neutrophil counts are positively predictive for each other when measured on the same day during cyclic chemotherapy for this smaller patient group.

The predictive relationship established by Kondo et al. (1999) was identified between monocytes with the onset of neutropenia counts 2 − 4 days prior to nadir. Unfortunately, due to the sampling rate in our data (monocytes and neutrophils measured once every 3 weeks), we were limited in our ability to discern a predictive relationship between monocytopenia and neutropenia. We therefore sought to leverage our model of granulo- and monocytopoiesis to understand the dynamics of monocyte and neutrophil production during cytotoxic chemotherapy with and without G-CSF support.

To predict monocyte and neutrophil dynamics during cyclic chemotherapy, we simulated chemotherapy administered every three weeks, similar to the DFCI ALL Consortium protocols described above. To quantify the effects of the treatment protocol, we calculated the time to nadir for both monocytes and neutrophils, as well as the beginning and duration of monocytopenic and neutropenic periods. Our results show that both the monocyte nadir and the beginning of monocytopenia precede the beginning of neutropenia, indicating that monocyte concentrations could be used to predict the onset of neutropenia (Figure 5). We find that the monocyte nadir robustly preceded the neutrophil nadir by at least three days in all cases, as in Kondo et al. (1999). As previously mentioned, this observation is perhaps to be expected, since monocytes and neutrophils share much of their development pathway; the lack of a mature monocyte reservoir may account for this time delay.

**Fig. 5.**
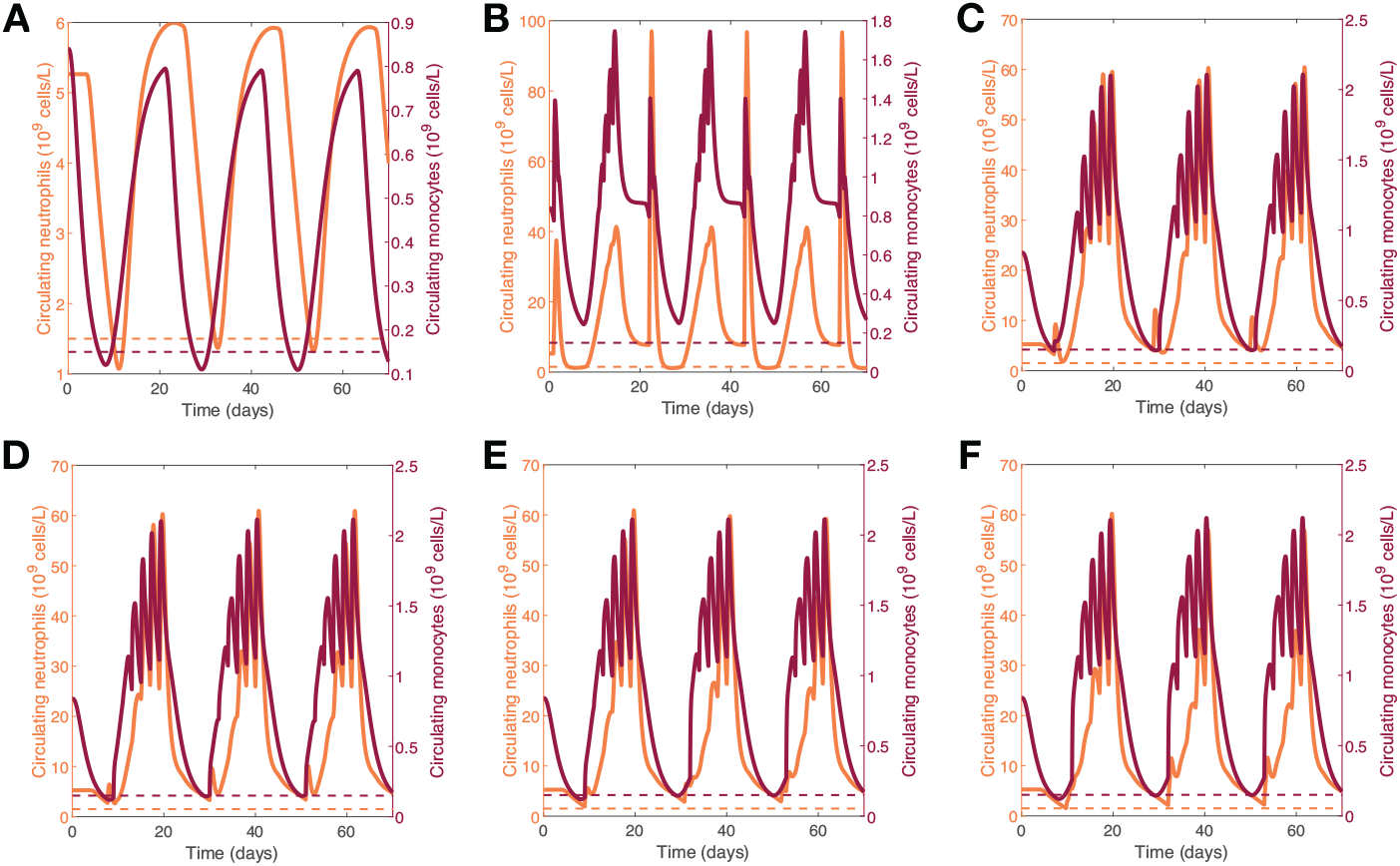
Modelling predicts that monocytopenia precedes neutropenia. A) Circulating monocyte and neutrophil concentrations in response to chemotherapy administered every 21 days with the thresholds for monocytopenia and neutropenia. B) Combination chemotherapy administered every 21 days with adjuvant G-CSF administered on 14 consecutive days beginning 1 day post chemotherapy. C) The results for therapy beginning immediately following the beginning of monocytopenia. D) Monocytopenia informed G-CSF schedule for therapy beginning one day following the beginning of monocytopenia. E) Monocytopenia informed G-CSF schedule for therapy beginning two days following the beginning of monocytopenia. F) Monocytopenia informed G-CSF schedule for therapy beginning three days following the beginning of monocytopenia. Solid dark purple: monocytes concentrations, dashed dark purple: monocytopenia threshold, solid light purple: neutrophils, dashed light purple: neutropenic threshold.

### 3.4 G-CSF support of the granulocyte-macrophage lineage during chemotherapy

G-CSF is used extensively during chemotherapy to avoid neutropenia. Accordingly, the dynamics of neutropenia during cyclic chemotherapy with and without G-CSF support have been extensively investigated (Mackey et al., 2020-submitted; Craig et al., 2015, 2016; Friberg et al., 2002; Quartino et al., 2014). Previous studies have suggested that delaying the administration of G-CSF after chemotherapy may improve the neutrophil response and help to mitigate ensuing neutropenia (Craig et al., 2015; Vainas et al., 2012; Krinner et al., 2013). However, monocyte dynamics during chemotherapy with G-CSF support are not as well-described. As such, we investigate the impact of G-CSF support of the granulocyte-macrophage lineage during chemotherapy, with a particular emphasis on its effects on monocytopenia to inform the use of G-CSF during cyclic cytotoxic chemotherapy.

We studied 3-week treatment cycles that each began with chemotherapy administered on day 1, followed by subcutaneous G-CSF administrations (Craig et al., 2016) on the *n*th day following the beginning of monocytopenia. As shown in Fig. 3, neutropenia consistently occurred three days after the onset of monocytopenia in the absence of G-CSF support. We therefore restricted the range of possible G-CSF administration days to be *n* = 0, 1, 2, 3 after the monocyte nadir. G-CSF administrations were then continued for *T* days until the end of the treatment cycle, and discontinued until day *n* in the subsequent cycle. The duration of neutropenia was the metric with which we measured the effectiveness of each schedule; schedules that minimized the neutropenic period were selected as being the most effective.

Our results show that neutropenia is completely mitigated by administering G-CSF every 2 days immediately following the onset of monocytopenia (Fig. 5). The same is true for G-CSF cycles of lengths *T* = 3, 4, 6, 7, 8. In the more physiologically realistic cases of G-CSF support beginning at least 1 day after the onset of monocytopenia (*n* = 1, 2, 3), administering G-CSF with a period of *T* = 2 completely avoids neutropenia.

Crucially, we find that *T* = 2 is not the only treatment frequency that eliminates neutropenia. Letting *T*_*i*_ denote a treatment schedule with G-CSF administered every *T* days beginning on day *i* following monocytopoiesis, we found that schedules where *T*_1_ = {3, 4, 6, 7, 8}, *T*_2_ = {4, 7, 8}, and *T*_3_ = {4, 8} also all eliminate neutropenia. This further supports previous modelling work that found that delaying the administration of G-CSF following chemotherapy may decrease the duration of neutropenia by avoiding emptying the mature marrow reservoir (Craig et al., 2016; Vainas et al., 2012; Krinner et al., 2013). Finally, our results suggest that it may be beneficial to only administer G-CSF once the anti-proliferative effects of cytotoxic chemotherapy are observed in the circulating monocyte concentrations, as monocytopenia precedes neutropenia. Interestingly, our results do *not* support the idea that the daily administration of G-CSF is an optimal therapeutic strategy.

## 4 Discussion

Despite the rapid progression in the development of biologics, cytotoxic chemotherapy remains the standard treatment for most cancers. Given its broad effects on not only cancerous growths but also hematopoietic cells, particular interest should be paid to quantifying blood cell dynamics during cytotoxic chemotherapy. To this end, mathematical modelling has played a significant role, helping to delineate the mechanisms of observed responses to chemotherapy and suggesting improved treatment strategies to help mitigate neutropenia while reducing the therapeutic burden to patients.

Here, to further the quantitative effort to characterize the toxic hematopoietic effects of cytotoxic chemotherapeutic approaches, we extended our previous model of granulopoiesis to include monocytopoiesis. For this, we developed a model of monocyte production beginning from hematopoietic stem cells to their evolution into non-classic monocytes in circulation. Model parameters were then obtained from existing literature sources and through parameter estimation. Parameter estimations were also cross-validated to different data to assess the accuracy of predictions. We also confirmed the robustness of our estimations through local sensitivity analysis.

Importantly, we used our model to characterize monocyte and neutrophil dynamics during chemotherapy in a clinical context. We first assessed the relationship between monocytes and neutrophil concentrations in close to 300 childhood ALL patients treated with 3-week cyclic chemotherapy, and found positive correlations in a subset of the total cohort. Unfortunately, we were unable to validate this observation in all patients.

Based on the results of Kondo et al. (1999), we were interested in quantifying the temporal cascade of monocytopenia and neutropenia, but were unable to assess this relationship due to the frequency of sampling in our data. We therefore simulated chemotherapy administered in 3-week cycles and predicted monocyte and neutrophil responses and found that monocytopenia preceded neutropenia by 3 days. We then investigated the effects of G-CSF administration concurrent to chemotherapy on the granulocyte-macrophage lineage, predicting that delaying the first administration of G-CSF administration until the onset of monocytopenia can reduce or completely eliminate neutropenia. While this prediction differs from the optimal G-CSF administration from our previous work (Craig et al., 2015), the onset of monocytopenia is a clinically relevant measurement.

Despite validating our parameter estimations and model predictions with previous studies, our approach is not without limitations. Some of the parameters used to simulate the monocytopoiesis model were either determined from experimental studies with a limited number of participants, or fit to data following the administration of exogenous G-CSF, which is not the primary regulator of monocyte production. In particular, the fitted maximal proliferation rate 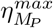 is very close to the (calculated) homeostatic proliferate rate 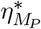. This is not surprising, as monocyte proliferation is thought to be independent of G-CSF, but illustrates the need to integrate other relevant cytokines involved in the control of monocytopoiesis when translating our model to infection and other inflammatory conditions. Moreover, the mathematical model has not been parameterized to specifically represent physiological differences present in patients with ALL. Accordingly, our modelling should be interpreted as a qualitative description of the hematopoietic response to chemotherapy, rather than a specific quantitative model. Nonetheless, our model was able to capture dynamics from a broad spectrum of studies in a variety of scenarios, and represents an important development for understanding the relationship between monocyte and neutrophils in the bone marrow and in circulation after cyclic chemotherapy with or without G-CSF support. This work therefore more broadly underlines the contribution of mathematical modelling to rationalizing clinical lines of investigation.

## Acknowledgements

MCM would like to thank Jim Murray for over 40 years of collegial friendship, and Prof. Dr. Klaus Pawelzik, Universität Bremen, Germany for his hospitality during the time this was written. All authors wish to thank Sanja Glisovic, Drs. Jean-Marie Leclerc, Yves Pastore, and Maja Krajinovic, and the patients enrolled in the ALL study at the CHU Sainte-Justine Research Centre.

